# Incomplete killing by antimicrobial metals generates transient antibiotic tolerance via delayed recovery in *Escherichia coli*

**DOI:** 10.64898/2026.06.19.733367

**Authors:** Merilin Rosenberg, Sandra Park, Angela Ivask, Niclas Nordholt, Frank Schreiber

**Affiliations:** Federal Institute for Materials Research and Testing (BAM), Berlin, Germany; Institute of Molecular and Cell Biology, University of Tartu, Tartu, Estonia

## Abstract

Incomplete killing is common after application of disinfectants or use of antimicrobial surfaces, leaving behind bacterial survivors with poorly characterized physiology. Here, we investigated the effects of short biocidal exposure to the common antimicrobial metals copper and silver as metal ions in suspension or under semi-dry conditions on solid metallic surfaces used against *Escherichia coli*. Survivors exhibited delayed regrowth and increased tolerance to bactericidal antibiotics without changes in MIC, consistent with a transient, non-genetic tolerance phenotype. When the magnitude of killing by silver or copper was matched between exposure formats, antibiotic tolerance patterns converged across metals and exposure conditions, indicating that the survivor state is governed more strongly by the severity of the primary lethal stress than by metal identity or exposure format alone. Incremental copper exposure revealed a dose-dependent increase in delayed regrowth and antibiotic tolerance but reduced tolerance to subsequent copper exposure. Post-exposure antibiotic tolerance as well as copper hypersensitivity phenotypes reverted after regrowth. Together, this supports a model of injury-linked delayed recovery rather than generalized stress priming. These findings demonstrate that incomplete disinfection can reshape survivor physiology, transiently increasing antibiotic tolerance through combined effects of selective survival acting on pre-existing phenotypic heterogeneity and acute injury-linked delayed recovery.

## Introduction

Biocides, including disinfectants and antimicrobial surface materials, are widely deployed to limit microbial transmission in healthcare, food-processing, and public environments. Their performance is typically evaluated as biocidal effect size under standardized conditions that require rapid multi-log reductions in viable counts (European Chemicals Agency., 2023; US Environmental Protection Agency, 2022). However, real-world conditions - characterized by variability in temperature, humidity, surface wetness, organic load, and inoculum properties - often result in incomplete killing leaving behind a substantial fraction of surviving cells (Campos et al., 2016; Cunliffe et al., 2021; Kaur et al., 2024). While the determinants of antimicrobial efficacy under such conditions are increasingly well characterized, the physiology of the survivors remains comparatively poorly understood.

Recent reviews further emphasize that the physiological consequences of biocide exposure cannot be inferred from nominal active-agent identity alone, because survivor outcomes depend on exposure conditions, mechanism of action, and the physiological state of the exposed cells (Maillard and Pascoe, 2024; O’Reilly et al., 2025). The resulting survivor states become important in the context of developing biocide tolerance and potential cross-resistance to antibiotics (Walczak et al., 2025). Most studies addressing interactions between biocides and response to antibiotics have focused on chronic and/or sublethal exposure regimes, during which bacteria adapt through regulated stress responses, genetic selection, or both (*e.g.* Fernandes et al., 2024; Nordholt et al., 2021). These and similar studies have provided important insights into mechanisms such as efflux-mediated cross-resistance, detoxification systems, and transcriptional reprogramming during growth (Kershaw, 2005; Liu et al., 2023; Yamamoto and Ishihama, 2005). However, disinfection events and fomite contacts in real-world settings are typically short, intense, and frequently occur under non-growth or multi-stress conditions. Under these constraints, inducible defense mechanisms may be temporally limited, suggesting that survival is more likely governed by pre-existing physiological heterogeneity and acute damage rather than fully deployed adaptive responses.

This distinction is particularly relevant in the context of antibiotic response. In antibiotic research, resistance refers to the heritable ability to grow at inhibitory concentrations, whereas tolerance reflects the ability to survive transient exposure to lethal concentrations without a change in the minimum inhibitory concentration, and persistence denotes survival of a subpopulation within a heterogeneous population (Balaban et al., 2019; Brauner et al., 2016). We acknowledge that there are differences in the fields of disinfection and antibiotic research regarding the somewhat confusing use of resistance and tolerance terminology (Kramer et al., 2026; Krewing et al., 2024; O’Reilly et al., 2025) that would complicate communicating research at the interface of the two fields. Therefore, in this study, tolerance is defined as the ability to survive transient exposures and resistance as the ability to grow at concentration above the MIC value of an antimicrobial agent. A key mechanism underlying increased tolerance to biocidal antibiotics is prolonged lag phase (delayed regrowth) (Fridman et al., 2014) originating from phenotypically heterogeneous populations (Hamill et al., 2020; Jõers and Tenson, 2016; Moreno-Gámez et al., 2020) or physiology induced by external factors (Hamill et al., 2020). Accordingly, colony appearance-time distributions (reflecting single cell lag-time distributions) provide a quantitative readout of recovery heterogeneity and a practical bridge between post-exposure physiology and antibiotic tolerance. Such tolerance-by-lag dynamics can arise from both regulated dormancy states and non-specific injury that prolongs recovery (Rotem et al., 2026). Short exposures to biocides may therefore generate antibiotic-tolerant survivor populations through a combination of selective survival of pre-existing phenotypic variants and induction of transient injury states.

Despite its potential importance, evidence for altered antibiotic tolerance following short biocidal exposures to biocides remains limited and fragmented. Observations across different agents, including copper, hypochlorite, and triclosan, suggest that such exposures can produce survivors with delayed regrowth or persistence-like phenotypes (An et al., 2026; Grey and Steck, 2001; Lin et al., 2017; Maertens et al., 2021; Westfall et al., 2019), but the related literature is scarce, experimentally heterogeneous and challenging to compare. In particular, it remains unclear to what extent post-exposure antibiotic tolerance reflects selection of pre-existing subpopulations, physiological changes induced by the biocide and/or the severity of the cellular damage.

Here, we tested whether short biocidal exposure to the widely used antimicrobial metals copper and silver generates survivor populations with altered post-exposure antibiotic tolerance. Copper and silver were selected because they are widely used antimicrobial metals with partially overlapping but context-dependent multi-target toxicity, allowing comparison of biocide-induced antibiotic tolerance across different biocidal effect sizes and exposure conditions, namely exposure to metal ions in suspension and to metallic surfaces under semi-dry conditions. Using *Escherichia coli* ATCC 8739, the widely used test strain for biocide efficacy testing as a model, we combined comparable biocidal exposures, lag-time distributions and post-exposure time-kill assays to test whether transient antibiotic tolerance arises predominantly from selection acting on pre-existing phenotypic heterogeneity, from injury-linked delayed recovery, or a combination of both.

## Materials and Methods

### Strains and media

All media, buffers and solutions were made with deionized water (MQ) and autoclaved at 121°C 15-20 min or filter-sterilized using syringe filters with 0.2 µm pore size. All bacterial manipulations were executed in aseptic conditions in biosafety cabinets. Media and buffers used are described in Table 1.

**Table 1.** Media and solutions.

| Media, buffers | Content |
| --- | --- |
| Luria-Bertani Broth (LB), liquid medium | 5 g/L yeast extract, 10 g/L tryptone, 5 g/L NaCl in L3022, Sigma-Aldrich®, Sigma-Aldrich Chemie GmbH |
| LB, solid medium | LB Broth with 15 g/L agar (A0949, AppliChem GmbH) |
| Nutrient Broth (NB) | 3 g/L meat extract (70164, Millipore®, Sigma-Aldrich Chemie GmbH), 10 g/L peptone (NCM0259B, Neogen), 5 g/L NaCl |
| Phosphate Buffered Saline (PBS) | 10-fold diluted stock of 8 g/L NaCl, 0.2 g/L KCl, 1.44 g/L Na<sub>2</sub>HPO<sub>4</sub>, 0.2 g/L KH<sub>2</sub>PO<sub>4</sub>; pH 7.1 |

*Escherichia coli* strain routinely used for efficacy testing of antibacterial agents was used in all experiments. *E. coli* DSM 1576 (ATCC 8739, NCIB 8545, WDCM 00012, Crooks strain) was ordered from the German Collection of Microorganisms and Cell Cultures GmbH. Here onwards, the more widely used synonymous ATCC 8739 strain number is used. 2^nd^ subculture of the received *E. coli* strain was inoculated into LB 0 broth, incubated for 24 h (180 rpm, 37°C), mixed 1:2 with 60% glycerol, aliquoted and stored at -80°C. For each experiment a new aliquot of the strain was used.

### Antimicrobial chemicals and surfaces

During antimicrobial exposures CuSO_4_ or AgNO_3_ as sources of copper and silver ions in liquid exposures or metallic copper and silver surfaces in semi-dry conditions were used. Round metal coupons with 20 mm diameter of 99% copper (Metroprint OY, Estonia) and 99.5% silver (Surepure Chemetals, USA) were used as antimicrobial surfaces.18×18 mm borosilicate microscopy cover slips (Corning Inc., USA) or sterile water were used as non-biocidal negative controls. All surfaces were rinsed with 70% ethanol and dried in aseptic conditions before exposures. Copper and silver surfaces were cleaned and reused for sustainable material use as described by Kaur et al. (Kaur et al., 2024). Experimental design with biocidal exposures, time-kill assays and lag time assessment is illustrated on Figure 1.

**Figure 1.**
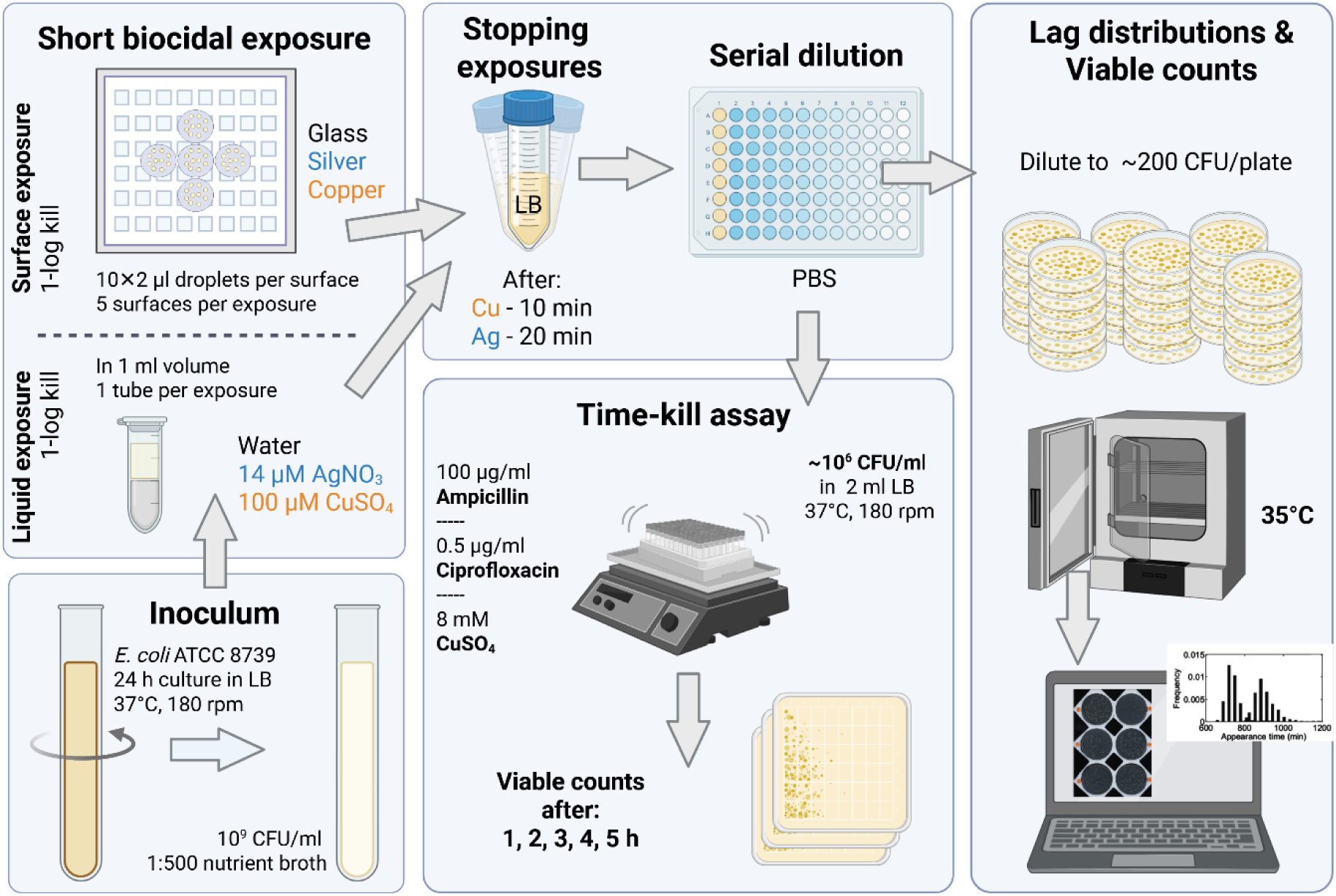
Experimental design. Short biocidal exposure workflow used to assess post-exposure recovery and tolerance phenotypes in *Escherichia coli*.

### Short biocidal exposure to copper or silver surfaces

24 h cultures of *E. coli* ATCC 8739 in LB were washed twice (5000 g, 10 min) with 1:250 diluted NB and photometrically adjusted to 2×10^9^ CFU/mL in the same medium. The suspension was diluted 1:1 with deionized water and 10×2 µL droplets (∼2×10^7^ CFU) were applied to each surface by means of an electronic stepper pipette. The inoculated surfaces were incubated in half-open Petri dishes in a biosafety cabinet at ambient conditions (21-24°C; 20-27% RH, cabinet lighting). Short biocidal exposures were tuned to produce incomplete disinfection (substantial but incomplete killing), enabling characterization of the survivor fraction. In the set of experiments depicted in Figure 2, bacteria were exposed to copper surfaces for 10 min and to silver surfaces for 20 min to achieve similar ∼1-log reduction in viable counts. Exposure time to glass control surfaces (10 or 20 min) was matched with exposure time to antimicrobial surface type. In the experiments depicted in Figure 3, bacteria were incrementally exposed to copper surfaces for 0.5 to 20 minutes coupled to 10 min control exposure on glass. Exposures were neutralized via transferring the inoculated surfaces into 10 mL fresh LB in 50 mL centrifuge tubes and briefly vortexed. This step minimizes continued killing after the defined exposure window by rapid dilution and complexation/speciation of metal ions in LB. Five surfaces (ma× 10^8^ CFU from control surfaces or ∼10^7^ CFU from 10 min biocidal exposures) were pooled into one neutralization tube after which the tube was vortexed at maximum intensity for 30 seconds. Bacterial suspensions in LB were immediately used for serial dilution in PBS for the following analyses. Any deviations from above description are mentioned in figure legends.

**Figure 2.**
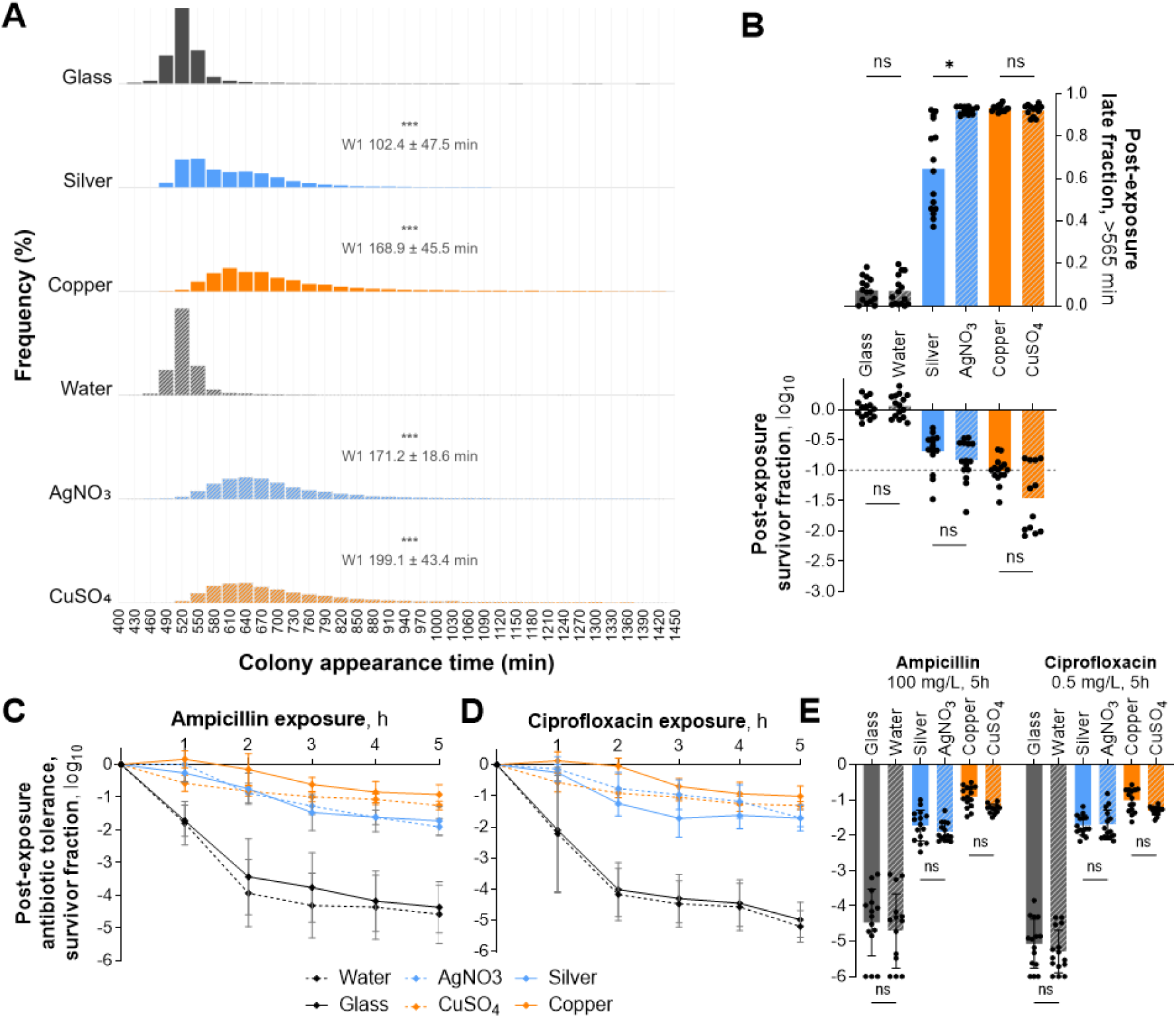
Short copper and silver exposure leads to survivors with delayed regrowth and increased antibiotic tolerance. Lag-time distributions (A), post-exposure survival and late fraction (B), antibiotic time-kill dynamics (C, D), and 5 h antibiotic tolerance (E) after short biocidal exposure to copper or silver in semi-dry surface (solid fill and lines) and liquid formats (patterned fill and dashed lines). Panel A shows lag-time distributions with significance codes indicating PERMANOVA comparisons against the respective glass or water controls, and Wasserstein-1 (W1 with ± SD) values summarize the average shift needed to align distributions with that respective control. Data from 5 independent experiments and 15 replicate exposures is presented (n=15, for CuSO_4_ n=12) as detailed in per-replicate data in Supplementary Fig S1. Paired liquid-versus-surface comparisons were performed at approximately matched killing for each metal after 10 min copper and 20 min silver exposures. Matched liquid-versus-surface exposure pairs were compared using non-parametric Kruskal–Wallis test with Dunn’s adjustment for multiple comparisons. For panels A, B and E, ns denotes P > 0.05, * P < 0.05, and *** P < 0.001.

**Figure 3.**
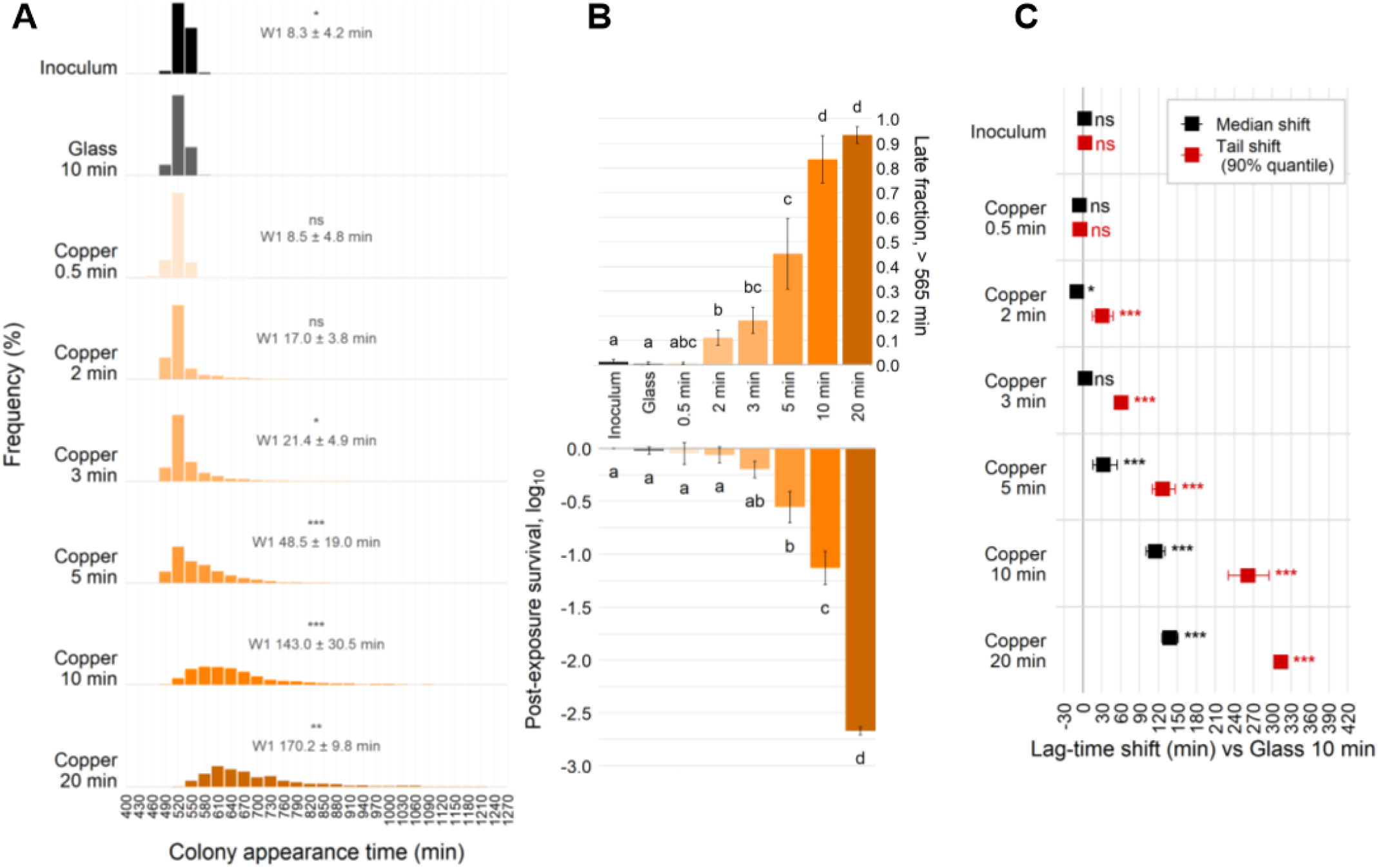
Regrowth delay increases with copper surface exposure severity. Lag-time distributions (A), post-exposure survival and late fraction (B), and lag-time shift estimates (C) after 0.5–20 min semi-dry exposure to copper surfaces. Panel A shows lag-time distributions with significance codes indicating PERMANOVA comparisons against the 10 min glass control, and Wasserstein-1 (W1 with ± SD) values summarize the average shift needed to align distributions with that control. Panel B shows decreasing viability together with increasing late fraction across the exposure series (±SD). Similarity groups were determined separately for survival and late fraction using Welch ANOVA followed by Games–Howell post-hoc comparisons at α = 0.05, with survival analyzed on the raw log_10_ scale and late fraction tested on the logit scale and displayed back on the raw 0–1 scale. Panel C shows bootstrap estimates of median and 90% quantile shifts relative to the 10 min glass control with ±95% CI error bars. For panels A and C, ns denotes P > 0.05, * P < 0.05, ** P < 0.01, *** P < 0.001, and **** P < 0.0001. Data from 3 independent experiments and 4-12 replicate exposures is presented as detailed in per-replicate data on Supplementary Fig S2. Supplementary Figures S8-10 provide complementary condition- and replicate-level and all-to-all distribution comparisons.

### Short biocidal exposure to CuSO_4_ or AgNO_3_ in liquid exposure medium

1 mM stock solutions of CuSO_4_ and AgNO_3_ in sterile water were prepared and stored in glass vials protected from light. The same stock solutions were used for all experiments. 24 h cultures of *E. coli* ATCC 8739 in LB were washed twice (5000 g, 10 min) with 1:250 diluted nutrient broth (NB) and photometrically adjusted to 2×10^9^ CFU/mL in the same medium. Metal salt solutions for the exposures were prepared immediately before exposures in twice the final exposure concentration. The bacterial suspensions were mixed 1:1 with 200 µM CuSO_4_ or 28 µM AgNO_3_ in final volume of 1 mL. Exposure times of 10 min for copper and 20 min for silver were used to achieve similar reduction in viable counts and match exposure time to surface exposures. Deionized water was used as a non-biocidal negative control instead of metal salt solution in liquid exposures. Exposure was neutralized via dilution and speciation of metal ions by transferring 1 ml of liquid exposure to 9 mL of fresh LB in a 50 mL tube and mixed by vortexing. Bacterial suspensions in LB were immediately used for serial dilution in PBS for the following analyses.

### Measurement of lag-time distributions

To acquire lag-time distributions of the survivors of short biocidal exposures, custom setup of the ScanLag system (Levin-Reisman et al., 2010) with 9 scanners in a 35°C microbiological incubator was used. 200 µL of appropriate serial dilutions of exposed bacteria expected to result in ∼200 colonies per plate were plated with sterile glass beads, covered by sterile dark fleece and loaded onto the scanners. After initial delay of 4 h, plates were scanned with 15 min intervals (69 h for data on Fig. 2 and 24 h for data on Fig. 3-5), depending on the experiment. Acquired time-series of images were analyzed for colony appearance time in MATLAB r2014a as described by Levin-Reisman et al. (Levin-Reisman et al., 2010). Number of individual experiments and exposure replicates included to and excluded from analysis are marked on Supplementary Figs. 1-2. Unless otherwise stated, individual exposures represent independent biological replicates, whereas multiple plates or image series derived from the same neutralized exposure were pooled to increase sensitivity. Replicates of exposures with ≥100 data points for colony appearance times across one to several imaged Petri dishes per survivor dilution were included in the downstream analysis due to lower counts possibly underestimating late fraction due to lower sensitivity.

### Measurement of antibiotic tolerance: time-kill assay

To assess the effect of copper and silver exposure on subsequent ampicillin and ciprofloxacin or CuSO_4_ tolerance, the survivors from short biocidal and control exposures were incubated in the presence of the bactericidal antibiotics ampicillin (Amp, 100 µg/mL) and ciprofloxacin (Cip, 0.5 µg/mL) or a bactericidal concentration of CuSO_4_ (8 mM) in 2 mL volume of fresh LB broth, 37°C,180 rpm. Bactericidal agents were added to the appropriate dilution of the exposed bacterial suspension in LB broth to achieve an estimated viable cell count of ∼10^6^ CFU/mL at the beginning of the time-kill assay. At predetermined time-points, an aliquot from each tube was serially diluted and drop-plated for viable counts. At the last time-point, remaining samples were concentrated by centrifugation (5000 g, 10 min) and plated for viable counts. Colonies were counted after 48 h of incubation at 37 °C.

### Controlling for emergence of genetic resistance

Stochastic emergence of stable resistance was assessed by an agar dilution MIC screen. For that 2 colonies from the last viable time point of antibiotic exposures in the time-kill assays were picked into 150 µL fresh LB in a 96-well plate, grown overnight (37 °C ;180 rpm), diluted 10^4^ times in PBS and spotted onto LB agar with respective antibiotics (0-4x the reference MIC: 8 mg/L Amp, 0.02 mg/L Cip) to detect stable changes in resistance. MIC values were scored for visible growth after 24 and 48 h incubation at 37 °C. As a common practice (Kowalska-Krochmal and Dudek-Wicher, 2021) spots with only 1-2 colonies at 1x MIC value were omitted due to technical variability at the threshold value.

### Stability of increased antibiotic tolerance and copper hypersensitivity after copper surface exposure

To test stability of tolerance phenotypes, bacteria were exposed to copper surfaces for 10 min as described above. 1 mL of the survivor suspension in LB was reinoculated into 9 mL of fresh LB medium and grown for 24 h at 37 °C, 180 rpm. Both before and immediately after the copper surface exposure as well as after overnight re-growth of the survivors, 5 h tolerance assay with 100 mg/L ampicillin or 8 mM CuSO₄ was conducted as described above for time-kill assay. Survivor fractions of the three sequential ampicillin or CuSO₄ exposures were compared to assess persistence or reversal of the tolerance phenotype.

### Data and statistical analysis

GraphPad Prism 10.4.1 was used for multiple comparisons and plotting of viability data. Downstream processing of lag-time data, distribution-level analyses, correlation/regression analyses, and plotting of the corresponding figures were performed in R (R Core Team, 2023), with ggplot2 (Wickham, 2016) for visualization. BioRender.com was used for schematic visualization. Unless otherwise stated, statistical tests were two-sided and assessed at α = 0.05. For figures presenting multiple comparisons, the applied ANOVA tests or its non-parametric analogues and multiplicity-corrected P-value adjustments are described in the respective figure legends. For association analyses, Pearson correlation was used to summarize linear associations and Spearman correlation to summarize monotonic associations. Because 5 h antibiotic-survival outcomes were left-censored at the detection limit, these outcomes were additionally modelled using Tobit regression (Lorimer and Kiermeier, 2007), allowing censored values to contribute information rather than being omitted. For incremental copper-exposure lag-distribution analyses, recovery kinetics were summarized by the median and upper lag-time quantiles together with the empirical late fraction, defined using a fixed control-derived threshold as the proportion of colonies appearing after 565 min, corresponding to the 99th percentile across untreated inoculum samples. This fixed threshold was used for enrichment-based analyses because it assigns the same absolute recovery-time boundary to pre- and post-exposure populations, whereas quantile-based summaries such as q90 were used as complementary threshold-independent descriptors of tail displacement. Complete, unbinned replicate-level lag-time distributions were compared using nonparametric pairwise energy distances (Rizzo and Székely, 2016), and condition effects on this distance matrix were tested by PERMANOVA (Anderson, 2001) with 9,999 permutations. This combination was used to compare whole lag-time distributions without assuming normality or a specific distributional shape due to potentially multimodal distributions as a result of phenotypic heterogeneity or post-exposure injury states. Distributional displacement effect size was alternatively summarized using the one-dimensional Wasserstein-1 (W1) distance, which can be interpreted as the mean absolute displacement in minutes required to optimally transport one lag-time distribution onto the other. Although transport- and energy-based distances are not standard in classical microbiology and lag-time analysis, related approaches have been applied to whole-distribution comparison in flow cytometry and single-cell biology, including Earth Mover’s/Wasserstein distance for comparing cytometric cell-population distributions and energy distance for testing global shifts in single-cell phenotypic distributions (Del Barrio et al., 2020; Orlova et al., 2016; Replogle et al., 2022; Schefzik et al., 2021). To localize where the displacement occurred, condition-wise differences in the median and upper-tail quantiles were estimated by non-parametric bootstrap resampling of replicate-level quantile summaries with 5,000 iterations. This framework was used because the lag-time distributions were asymmetric with potentially a second late-lag peak and tail behavior was biologically central to the study question. Thus, energy distances with permutation testing asked whether the distributions were globally different, W1 summarized the overall one-dimensional effect size, bootstrap-based median and tail shifts localized the change to the center or right tail of the recovery distribution, and late fraction followed the biological shift across a fixed threshold value established based on untreated controls. Where multiple pairwise or matrix-wise comparisons were made, P values were adjusted using the Benjamini–Hochberg false-discovery-rate procedure.

## Results

### Short exposure to copper and silver generates survivors with delayed regrowth and increased antibiotic tolerance

To assess the immediate physiological consequences of incomplete disinfection on antimicrobial surfaces, stationary-phase *Escherichia coli* ATCC8739 cells were exposed to copper and silver surfaces under semi-dry conditions or respective metal salts in suspension adjusted to produce approximately 1-log reduction in viable counts (Fig. 2B). To match the killing magnitude, 10 min exposure to copper and 20 min exposure to silver were used. Across all exposure formats, surviving populations displayed right-shifted lag-time distributions relative to non-biocidal controls, indicating generally delayed regrowth (Fig. 2A) and expansion of late fraction (Fig. 2B).

Antibiotic time-kill assays after exposure to copper or silver revealed biphasic killing dynamics for both ampicillin and ciprofloxacin (Fig. 2C, D), consistent with the presence of subpopulations exhibiting increased survival during prolonged antibiotic exposure. Despite differences in nominal antimicrobial potency between copper and silver and between surface and liquid exposure formats (Fig 2B), post-exposure antibiotic tolerance levels were broadly similar for both metals and exposure formats (Fig. 2E) when killing magnitude was matched (Fig. 2B). Minimum inhibitory concentrations remained unchanged across all conditions (Supplementary Table S1), indicating that the observed phenotype reflects altered antibiotic tolerance rather than resistance. These results show that short biocidal metal exposures can generate survivor populations with delayed recovery and increased antibiotic tolerance. Thus, despite different metals and exposure conditions, copper and silver exposures with comparable biocidal effect size converge on a similar post-exposure phenotype, supporting biocidal effect size rather than agent identity as the dominant determinant of survivor physiology.

Semi-dry exposure to model antimicrobial copper surfaces demonstrated the lowest variation in post-exposure survival and lag-time distributions among the different conditions examined (Fig. 2B; Supplementary Fig. S1) and was selected for the following incremental exposure series used for mechanistic analysis.

### Regrowth delay and antibiotic tolerance increase with copper surface exposure severity

To examine the relationship between copper surface exposure severity and post-exposure survivor phenotype, bacteria were subjected to incremental copper surface exposures ranging from non-lethal 0.5 min to strongly biocidal 20 min exposure. Due to the dynamic changes in metal ion release, speciation and inoculum volume changes in semi dry surface exposure format as well as both release-based and contact-killing properties of copper surfaces, incremental exposure time was used to control the exposure dose and respective biocidal effect size. Time-controlled biocidal effect size was used as a population-level proxy for exposure severity among the surviving populations as viable counts decreased with increasing exposure duration (Fig. 3B). Increasing exposure severity (Fig. 3B) resulted in a progressive rightward shift in lag-time distributions (Fig. 3A, Supplementary Fig. S2), with changes primarily driven by expansion of the late fraction (Fig. 3C) as the tail-shift preceded the median shift. Post-exposure antibiotic tolerance also increased with copper surface exposure severity, as indicated by higher survival during ampicillin and ciprofloxacin time-kill assays (Fig. 4A, B). At the replicate level, strong negative correlations were observed between post-exposure late fraction and both survival and antibiotic tolerance, while late fraction was positively associated with antibiotic tolerance (Fig. 4C–E). These relationships were robust to the use of alternative lag-time metrics, including the 90% quantile (Supplementary Fig. S3). Thus, the fixed >565 min late-fraction threshold provided a biologically anchored readout of entry into the control-defined late-recovery regime, while q90 confirmed that the association between delayed recovery and antibiotic tolerance was not an artefact of that single threshold. Together, these results indicate that increasing biocidal exposure severity is associated with delayed recovery and increased antibiotic tolerance among surviving populations.

**Figure 4.**
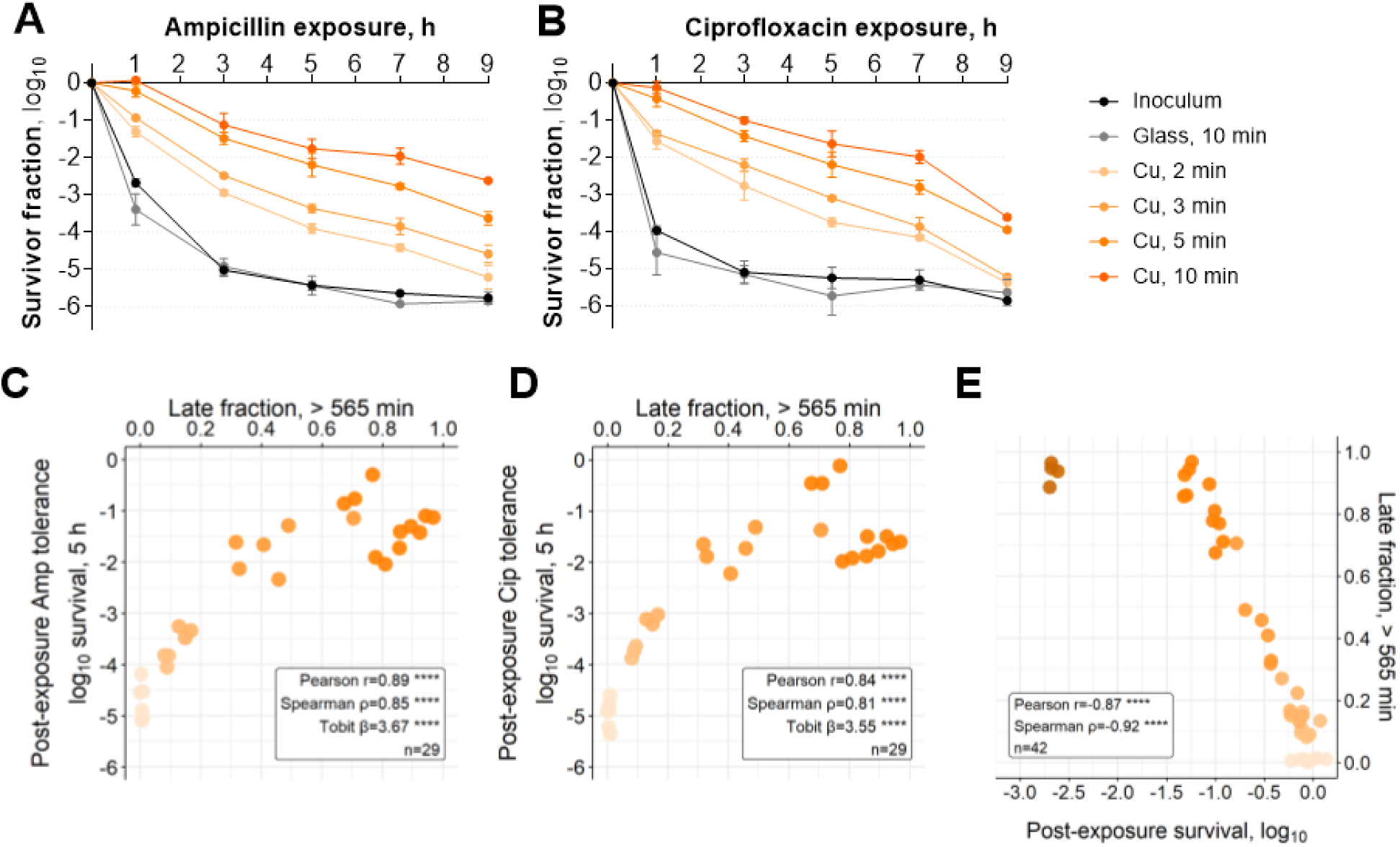
Antibiotic tolerance increases with delayed recovery after incremental copper surface exposure. Time-kill dynamics of ampicillin (A) and ciprofloxacin (B) after incremental semi-dry copper surface exposures, together with replicate-level associations between late fraction and 5 h antibiotic survival (C, D) or survival after copper surface exposure (E). Full time-kill assays with antibiotics on panels A and B were performed after 2–10 min copper exposures as 0.5 min (beige) and 20 min (brown) exposures yielded either no killing or too few survivors for time-kill experiments, respectively. Points in the association plots are gradient-colored by copper exposure duration. Pearson (linear) and Spearman (non-linear monotonic) correlation, and Tobit regression (where antibiotic-survival values are left-censored at the detection limit) coefficients are shown to summarize associations. Data from 3 independent experiments and 4-12 replicate exposures originating from Fig. 3, as detailed on Supplementary Fig. S2. **** denotes P < 0.0001 for the association. Corresponding analyses using the 90% lag-time quantile in addition to late fraction are shown in Supplementary Fig. S3.

**Figure 5.**
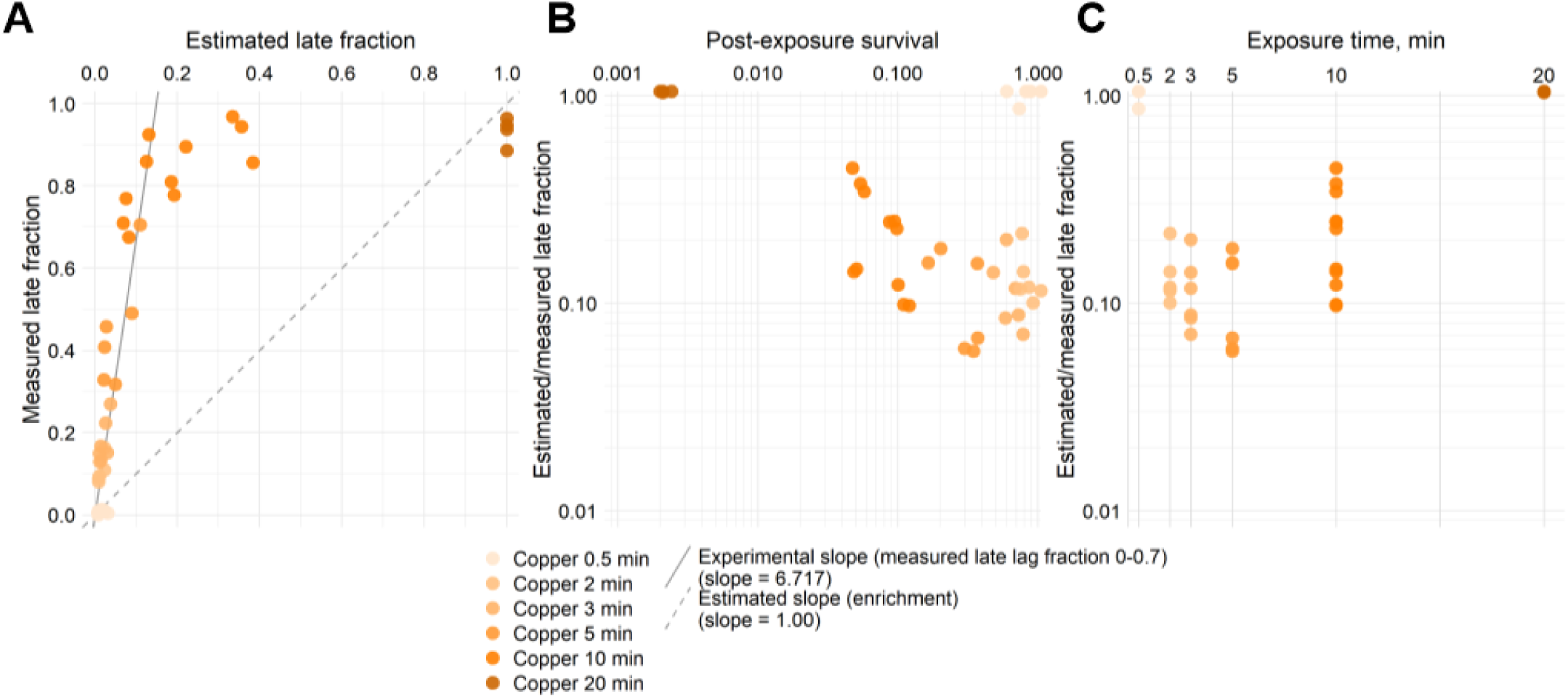
Delayed regrowth after copper surface exposure exceeds the enrichment-only late fraction expectation. Comparison of the measured post-exposure late fraction (p_post_) with the enrichment-only prediction min(1, p_0_/x), where p_0_ is the pre-exposure late fraction and x is post-exposure survival (data from Fig. 3). Because both p₀ and p_post_ were calculated using the same fixed >565 min control-derived threshold, the comparison tests enrichment of cells that would fall into the late-recovery tail of the untreated population. Panel A plots the measured late fraction against the enrichment-only expectation. The dashed diagonal indicates equality expected if the post-exposure phenotype were explained solely by selective survival of a pre-existing delayed subpopulation. Systematic deviations above the diagonal therefore indicate excess delay beyond the enrichment-only expectation. Panels B and C show the ratio between the enrichment-only expectation and the measured late fraction, min(1, p_0_/x)/p_post_, as a function of post-exposure survival and copper exposure time, respectively. Values below 1 indicate that the measured late fraction exceeds the enrichment-only prediction. Exposures in which survival is equal to or smaller than the pre-exposure late fraction (x ≤ p_0_) cluster below the diagonal in panel A because the enrichment-only estimate approaches its upper bound of 1 by construction. This may reflect slight underestimation of the selectively surviving subpopulation by the empirical >565 min threshold, overlap of the left tail of the delayed subpopulation with the main population, or incomplete selective survival in which some short-lag cells survive and some long-lag cells die. At low survival (x ≤ p0), the enrichment-only estimate is therefore not by itself strongly diagnostic.

### Delayed regrowth after copper surface exposure cannot be explained by only selective survival of a pre-existing late-lag subpopulation

Across the exposures shown in Figures 2–4, we observed experiment-to-experiment variation in the baseline late fraction and in the associated antibiotic tolerance. This variation likely reflects differences in the pre-existing late-cell fractions of individual 24 h tube-grown liquid cultures, as illustrated in Supplementary Fig. S4. Furthermore, additional experiments with cultures from different growth conditions than those used in the main experiments (Fig. 2-5) showed that *E. coli* can have inherently larger late fractions that result in higher antibiotic tolerance (Supplementary Fig. S4). For example, 72 h tube-grown cultures (Supplementary Fig. S5) and 24 h flask-grown cultures (Supplementary Fig. S6) with distinctive late fraction peaks were inherently more tolerant to antibiotics than the 24 h tube-grown cultures. The 72 h cultures with larger pre-existing late fraction showed increased tolerance to both copper surfaces and antibiotics compared with otherwise identically cultivated 24 h cultures (Supplementary Fig. S5).

To test whether the observed increase in delayed regrowth after incremental copper surface exposure (Fig. 3) could be explained solely by selective survival of a pre-existing long-lag subpopulation, the measured post-exposure late fraction was compared to an enrichment-only expectation derived from per-replicate pre-exposure late fraction and copper survival (Fig. 5A). In case of selective survival of a long-lag subpopulation, post-exposure late fraction would be proportional to survival. Across the incremental copper surface exposure series, the observed late fraction systematically exceeded this expectation, indicating that selective survival alone is insufficient to account for the magnitude of the observed lag shift.

Deviation from the enrichment-only prediction was evident already at short exposure times (Fig. 5C), with no significant killing (Fig. 5B and 3B) suggesting that the increase in delayed regrowth does not require extensive elimination of the short-lag population. These results indicate that additional processes beyond selective survival contribute to the emergence of delayed recovery phenotypes following copper exposure.

### Post-exposure phenotype demonstrates divergent but reversible antibiotic and copper tolerance

To further characterize the nature of the survivor state after copper surface exposures, tolerance to subsequent copper exposure of cultures with different initial late fraction sizes was assessed. In contrast to the increased tolerance observed for antibiotics, survivors of copper surface exposure exhibited reduced tolerance to subsequent CuSO₄ challenge (Supplementary Fig. S5). While the larger pre-existing late-fraction of 72 h cultures increased survival on copper surfaces and during antibiotic exposure compared to 24 h cultures, the larger initial late fraction did not protect bacteria from CuSO₄ challenge after copper surface exposure (Supplementary Fig. S5). The latter indicates that CuSO_4_ hypersensitivity after copper surface exposure is caused by the preceding toxic exposure and not directly by the inherent properties of the untreated culture.

Both the antibiotic tolerance and copper hypersensitivity phenotypes after copper surface exposure were transient and reverted after regrowth to stationary phase, with post-regrowth populations displaying ampicillin and CuSO_4_ tolerance comparable to untreated controls (Supplementary Fig. S7). These observations indicate that short copper exposure generates a reversible physiological state characterized by delayed recovery and altered tolerance properties.

## Discussion

Our results support a framework in which incomplete killing by short exposures to copper and silver generates survivor populations with transiently altered physiology characterized by delayed recovery and increased antibiotic tolerance. Across both copper and silver exposures, and across semi-dry surface and liquid exposure formats, antibiotic tolerance patterns were primarily determined by the severity of the biocidal insult rather than by metal identity or exposure format. This convergence suggests that different antimicrobial agents can drive cells into functionally similar recovery states when the extent of damage is comparable. Importantly, this finding implies that the physiological consequences of incomplete killing by *e.g.* disinfectants or antimicrobial surfaces may be predictable based on biocidal effect size rather than agent-specific properties alone.

A central question addressed in this study is whether post-exposure tolerance arises predominantly from selective survival of pre-existing phenotypic variants, from *de novo* physiological changes induced by copper and silver such as acute injury or phenotypic reprogramming or a combination of both selective survival and metal-induced changes. The incremental copper exposure time series provided evidence that selective survival of a pre-existing late-lag subpopulation alone was insufficient to explain the observed phenotype. While regrowth delay increased with copper exposure severity, the resulting late fraction was not proportional to copper survival and this deviation emerged already at low levels of killing. On the other hand, short biocidal exposure times make mechanisms requiring substantial *de novo* transcriptional reprogramming, protein synthesis and accumulation less likely to be the primary drivers during the defined exposure window. Instead, the results support rapid transitions into altered physiological states as a consequence of acute injury as one of the main drivers for increased antibiotic tolerance. These findings are consistent with the broader framework of phenotypic heterogeneity, in which pre-existing variability interacts with environmental stress to shape survival outcomes (Balaban et al., 2004; Nordholt et al., 2024). These results are also conceptually consistent with the distinction between regulated and disrupted growth arrest due to which the same stress, applied either abruptly or gradually, can lead to fundamentally different recovery dynamics (Kaplan et al., 2021) and similarly, elevated antibiotic tolerance can arise from physiologically distinct states (Rotem et al., 2026).

The combined phenotype of increased tolerance to bactericidal antibiotics but decreased tolerance to subsequent copper exposure further constrains the underlying mechanism. A generalized stress-priming model would predict cross-protection across stressors, whereas the observed divergence indicates that the post-exposure state is not broadly protective. Instead, this pattern is consistent with a condition that reduces susceptibility to growth-associated antibiotic killing by ampicillin (Tuomanen et al., 1986; Varik et al., 2016) and ciprofloxacin (Sampaio et al., 2022; Smirnova and Oktyabrsky, 2018) while increasing vulnerability to repeated metal exposure. This pattern is consistent with a recovery state following acute injury affecting core cellular processes such as membrane integrity, redox homeostasis, and macromolecular stability. Injury that does not lead to loss of viability can affect recovery kinetics and culturability. Delayed resumption of growth has been proposed as one consequence of microbial injury (Shao et al., 2023; Tsuchido, 2023; Wesche et al., 2009). Metal-induced oxidative stress, protein damage, and disruption of cellular homeostasis are well-established effects of copper and silver (Grass et al., 2011; Lemire et al., 2013; Liu et al., 2023) and may prolong recovery without conferring protection against additional stress. Such injury-associated states may transiently reduce metabolic activity and growth rates, thereby decreasing susceptibility to antibiotics that rely on active cellular processes.

Within this framework, delayed regrowth emerges as a unifying driver of antibiotic tolerance, regardless of whether it originates from pre-existing heterogeneity or injury-induced growth arrest. Our data show that both intrinsic variation in lag-time distributions resulting from different culture conditions as well as exposure-induced shifts toward longer recovery times are associated with increased antibiotic tolerance. This supports the concept of tolerance-by-lag (Fridman et al., 2014) and indicates that incomplete killing acts on a pre-structured population while simultaneously reshaping it through damage-induced physiological changes. The transient nature of the tolerance phenotype further supports a non-genetic mechanism. The rapid reversion after regrowth to stationary phase argues against stable epigenetic or bistable regulatory states (Scanlon et al., 2025) as dominant contributors and instead favors reversible physiological perturbations that are consistent with reversal of phenotypic memory in stationary phase (Faigenbaum-Romm et al., 2025).

From an applied perspective, this transient window of tolerance after short biocidal exposure may nevertheless be biologically relevant, as it could overlap with transmission events or early stages of infection. Increased tolerance during this window may allow a subset of cells to survive initial treatment, potentially facilitating subsequent adaptation or resistance evolution under continued selection (Levin-Reisman et al., 2017; Windels et al., 2019). These findings have direct implications for infection control and antimicrobial stewardship. This is particularly relevant in healthcare settings, where exposure to partially disinfected surfaces, antimicrobial surfaces or antimicrobially coated medical devices may precede or coincide with antibiotic therapy. Although such effects do not constitute resistance to antibiotics, they may influence treatment efficacy and contribute to persistence of infections (Walczak et al., 2025). More broadly, these results highlight that antimicrobial interventions can shape not only population size but also physiological cell states, with downstream consequences for susceptibility to subsequent treatments.

Several limitations should be considered when generalizing the results of this study. The study focuses on a single *E. coli* strain and widely used antimicrobial metals under defined conditions, and the extent to which these findings translate across clinically relevant pathogens, including opportunistic and multidrug-resistant species as well as other biocide classes, remains to be determined. In addition, the mechanistic basis of the injury state is inferred from phenotypic readouts rather than directly measured. In cases of incomplete killing, phenotypic screens and multi-omics methods face severe challenges in distinguishing between signals from the majority of dead cells, delayed post-exposure killing of irreversibly damaged viable cells (Fanous et al., 2025) and true survivors. Future work integrating single-cell approaches, metabolic and redox reporters, and molecular assays of damage and repair processes would help to resolve the underlying mechanisms more precisely and to determine how broadly these dynamics apply across organisms and environments.

Current academic and standardized evaluations of antimicrobial materials primarily emphasize efficacy endpoints, such as log-reductions in viable counts under defined test conditions. Our results suggest that, in cases of incomplete killing, these quantitative endpoints may not fully capture application-relevant outcomes, because the surviving fraction can differ physiologically from the starting population. Incorporating selected post-exposure readouts of survivor physiology, including delayed recovery and altered tolerance to subsequent antimicrobial exposures, could therefore complement conventional efficacy testing and improve the predictive value of laboratory assays for downstream risks such as biocide tolerance and antibiotic cross-tolerance.

## Conclusion

In *E. coli* ATCC 8739, short biocidal copper or silver exposures that model incomplete disinfection leave behind survivors in a transient physiological state characterized by delayed regrowth and increased tolerance to bactericidal antibiotics without a corresponding MIC shift. The physiological state of the survivor cells with respect to delayed regrowth and antibiotic tolerance was more closely related to the magnitude of the primary biocidal effect size as compared to metal identity or exposure conditions (semi-dry or liquid).

Increased copper surface exposure severity led to transiently increased regrowth delay and antibiotic tolerance and inversely, decreased tolerance to additional copper stress. At the same time, increasingly delayed regrowth was not proportional to copper survival. Therefore, selective survival of pre-existing late-lag subpopulation was insufficient to fully explain the increased post-exposure late fraction and the associated antibiotic tolerance. Together, this is consistent with a model that favors injury-linked delayed recovery over generalized stress priming as a mechanism that explains the observed phenotype.

More broadly, these results indicate that incomplete disinfection can reshape survivor physiology through combined effects of acute injury and selective survival acting on pre-existing phenotypic heterogeneity, and that post-exposure physiology should be considered alongside killing efficacy when designing and evaluating antimicrobial interventions and their downstream consequences.

## Acknowledgements

This research received funding from, Estonian Research Council (PRG1496), European Union Twinning project FAST-Real (101159721) and from the Estonian Ministry of Education and Research under projects TK210 and TEM-TA55. Academic mobility of MR and SP was supported by FEMS Research & Training grant, Kristjan Jaak National Scholarship Program (funded by the Ministry of Education and Research of the Republic of Estonia), Estonian Doctoral School (University of Tartu project “Cooperation between universities to promote doctoral studies” (2021–2027.4.04.24-0003)).

## Supplementary Figures and Tables

**Supplementary Table S1.** Antibiotic susceptibility changes after primary biocidal or control exposure and subsequent ampicillin or ciprofloxacin exposure across exposures presented on Figures 1-3. Number of isolated colonies yielding growth on LB agar with or without antibiotic is presented. Cultures derived from single colonies picked from the last antibiotic exposure time points with observed survivors in antibiotic time-kill experiments (Fig. 2 and 4) all yielded confluent growth on LB agar with no antibiotic and no growth at 1-4x MIC. No true inheritable antibiotic resistance was observed after antibiotic time-kill assays.

|  |  | Ampicillin, mg/L |  |  | Ciprofloxacin, mg/L |  |  |  |  |
| --- | --- | --- | --- | --- | --- | --- | --- | --- | --- |
|  |  | 0 | 8 | 16 | 32 | 0 | 0.02 | 0.04 | 0.48 |
|  |  |  | 1x MIC | 2x MIC | 4x MIC |  | 1x MIC | 2x MIC | 4x MIC |
| Unexposed culture |  | 6 | 0 | 0 | 0 | 6 | 0 | 0 | 0 |
| Control, liquid exposure, 10-20 min |  | 15 | 0 | 0 | 0 | 15 | 0 | 0 | 0 |
| Control, surface exposure, 10-20 min |  | 23 | 0 | 0 | 0 | 23 | 0 | 0 | 0 |
| Copper surface, 10 min |  | 26 | 0 | 0 | 0 | 26 | 0 | 0 | 0 |
| Silver surface, 20 min |  | 15 | 0 | 0 | 0 | 15 | 0 | 0 | 0 |
| CuSO<sub>4</sub> in suspension, 10 min |  | 12 | 0 | 0 | 0 | 12 | 0 | 0 | 0 |
| AgNO<sub>3</sub> in suspension, 20 min |  | 15 | 0 | 0 | 0 | 15 | 0 | 0 | 0 |

---

**Supplementary Figure S1.**
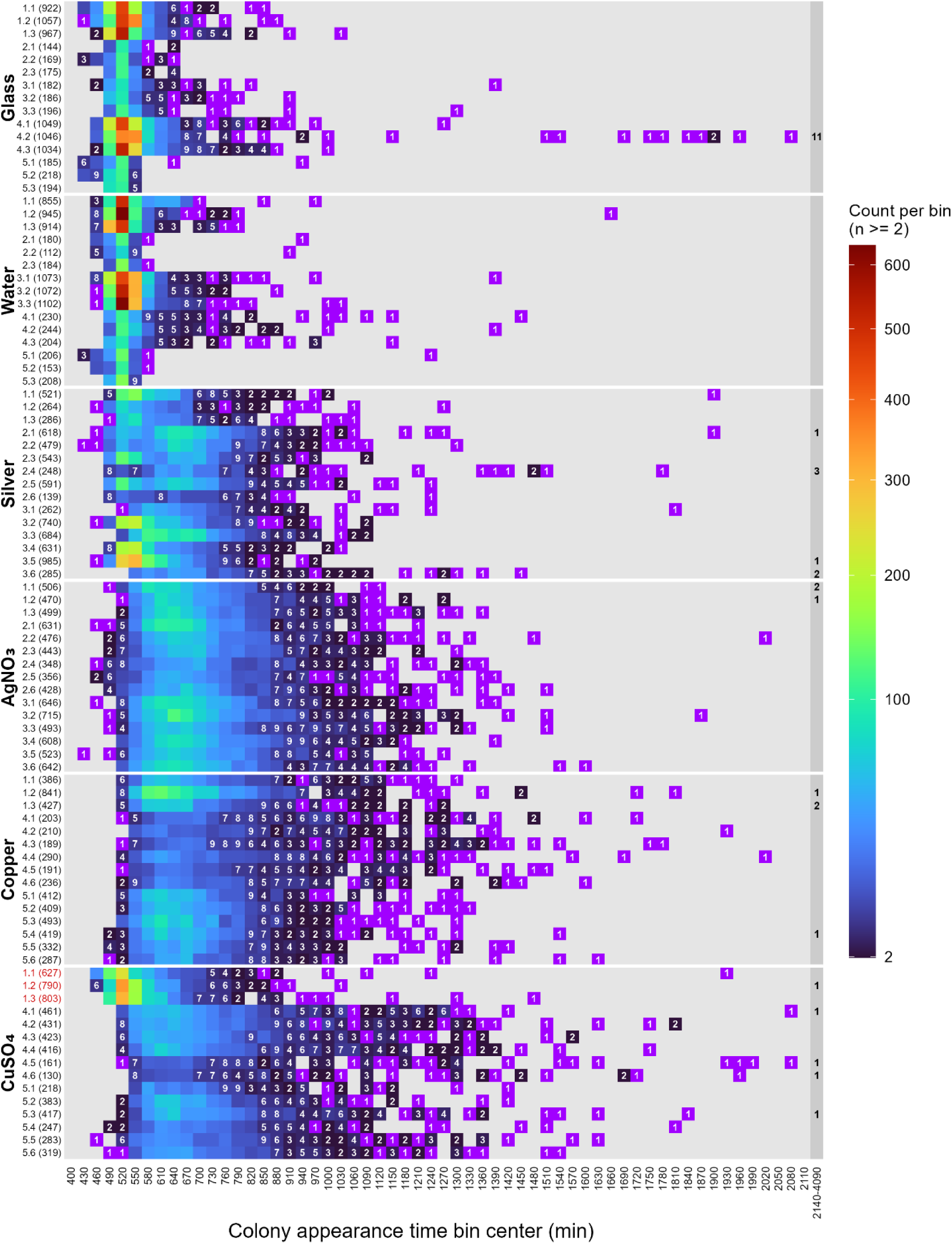
Individual lag-time distributions after biocidal or control exposures from 5 experiments underlying the averaged results in Fig. 2. Number of detected colonies across all imaged Petri dishes per replicate is marked in brackets next to each replicate exposure marked as “experiment.replicate(colonies)”. Bins with single detected colonies are highlighted in purple for visual contrast and single detected colonies from bins 2140-4090 pooled to last summary bin on grey background. Replicates removed from analysis and figures due to inconsistent results in viability and lag-time distributions are marked in red. The affected CuSO_4_ exposures 1.1-1.3 id not result in sufficient biocidal effect size reflected also in the unshifted lag-time distributions. AgNO_3_ exposures produced relatively consistent lag-time distributions, whereas silver surface exposure showed substantial inter- and intra-experiment variability. This pattern is consistent with variable drying of individual inoculum droplets and local microenvironmental conditions during the 20 min semi-dry surface exposure. As silver loses antibacterial activity after drying (Kaur et al., 2024), that can produce mixed post-exposure populations experiencing different effective silver stress. Lacking right tails of replicates 2^nd^ and 5^th^ experiment in case of glass and water control exposures possibly reflect inoculum properties further detailed on Supplementary Fig. S4.

**Supplementary Figure S2.**
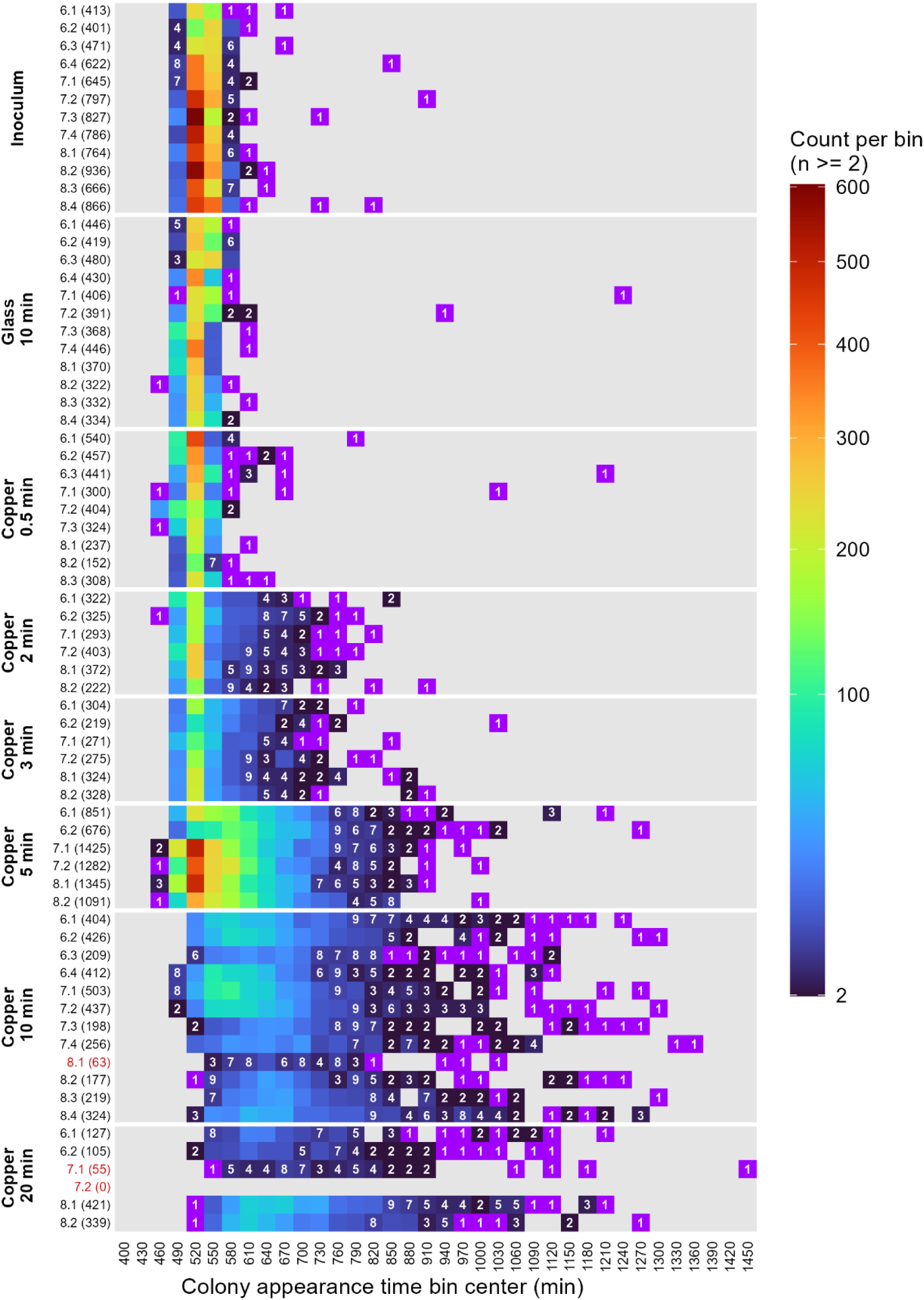
Individual lag-time distributions before and after 0.5-20 min copper surface or 10 min glass control exposures underlying the averaged results in Fig. 3. Data from 3 experiments with 2-4 replicate exposures is presented. Number of detected colonies across all imaged Petri dishes per replicate is marked in brackets next to each replicate exposure marked as “experiment.replicate (colonies)”. Bins with single detected colonies are highlighted in purple for visual contrast. Replicate exposures removed from analysis and figures due to ≤100 colonies per replicate after higher than expected biocidal effect size are marked in red.

**Supplementary Figure S3.**
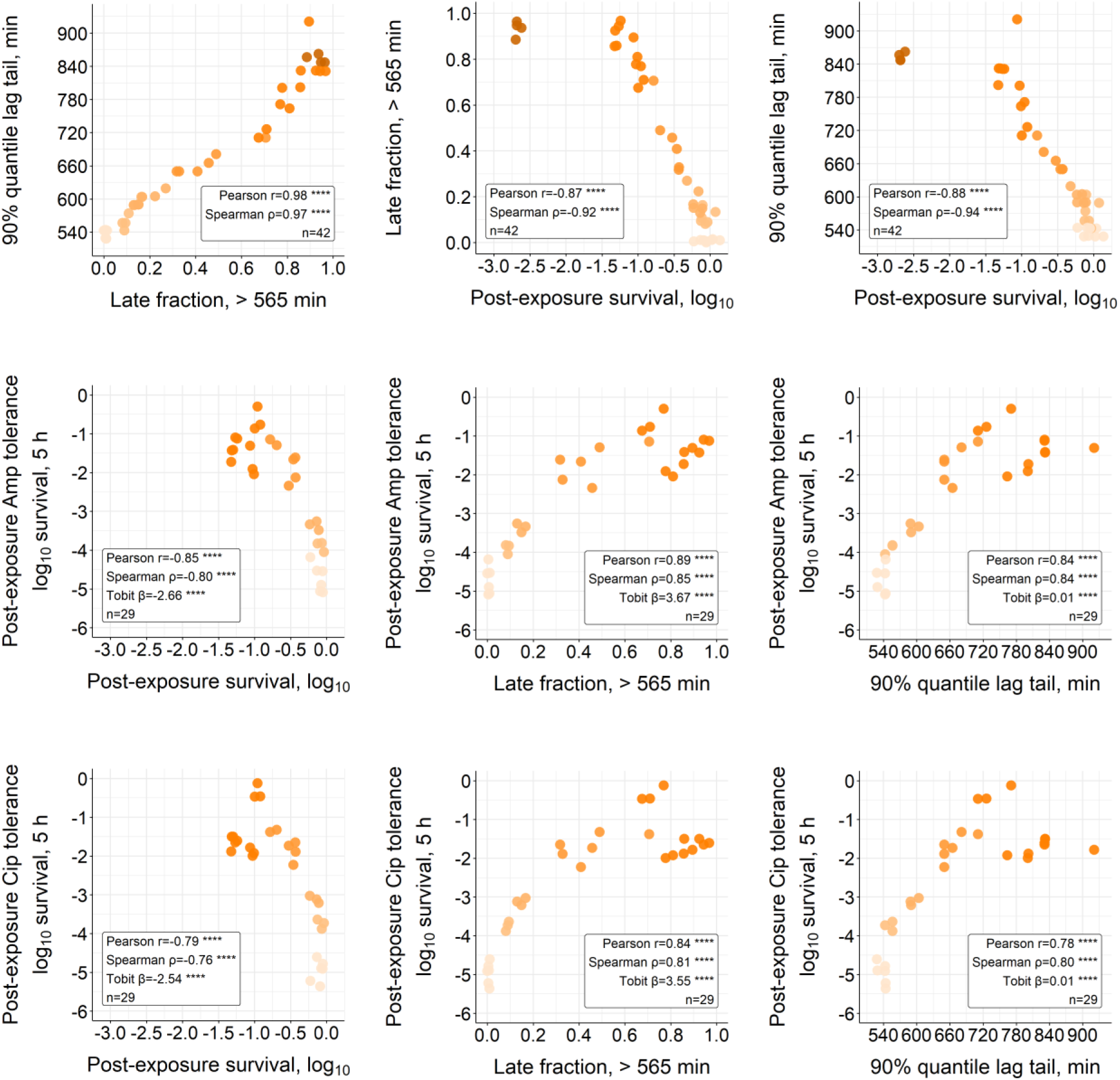
Copper-only replicate-level association plots relating post-exposure survival and antibiotic tolerance to delayed regrowth metrics in the incremental copper exposure series. The upper row shows post-exposure survival versus late fraction and versus the 90% lag-time quantile. The middle row shows post-exposure ampicillin tolerance (5 h, log_10_ survival) versus post-exposure survival, late fraction, and the 90% lag-time quantile. The lower row shows post-exposure ciprofloxacin tolerance versus the same variables. Each point represents one exposure replicate and is gradient-colored by copper exposure duration (from 0.5 min beige to 20 min brown). Insets report Pearson and Spearman correlation coefficients, and Tobit regression coefficients are shown where antibiotic-survival values are left-censored at the colony-counting detection limit. These panels complement Fig. 4C–E by showing that the q90 lag tail captures the same overall pattern as late fraction: later recovery tails are associated with higher antibiotic tolerance and lower post-exposure survival.

**Supplementary Figure S4.**
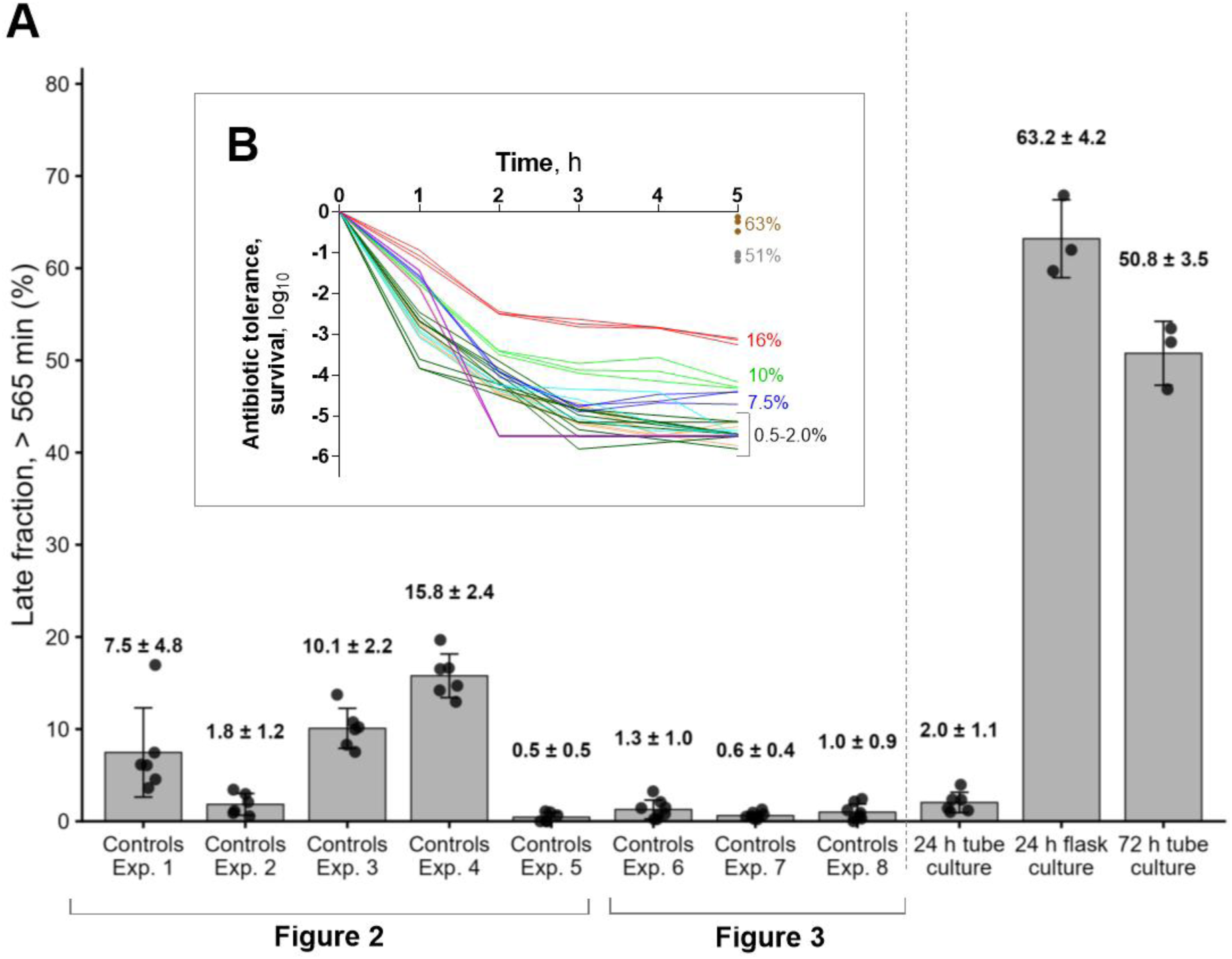
Baseline late fraction is associated with baseline ampicillin tolerance. Late fraction (>565 min) in untreated controls from the experiments shown in Figs. 2 & S1 (glass and water exposures) and Fig. 3 & S2 (inoculums and glass exposures) and in selected comparison cultures with inherently different late fractions from Figs. S5 and S6 (A), together with corresponding replicate-level ampicillin time-kill profiles color-coded for control/initial late fraction (B). Inoculums prepared from 24 h tube cultures and subjected to non-biocidal exposures contained variable late fractions from <1% to 16% and resulted in inter-experiment variation in baseline ampicillin tolerance. The figure therefore illustrates that inoculum-to-inoculum physiological variation is a likely contributor to interexperiment variability in both post-exposure lag distributions and antibiotic tolerance.

**Supplementary Figure S5.**
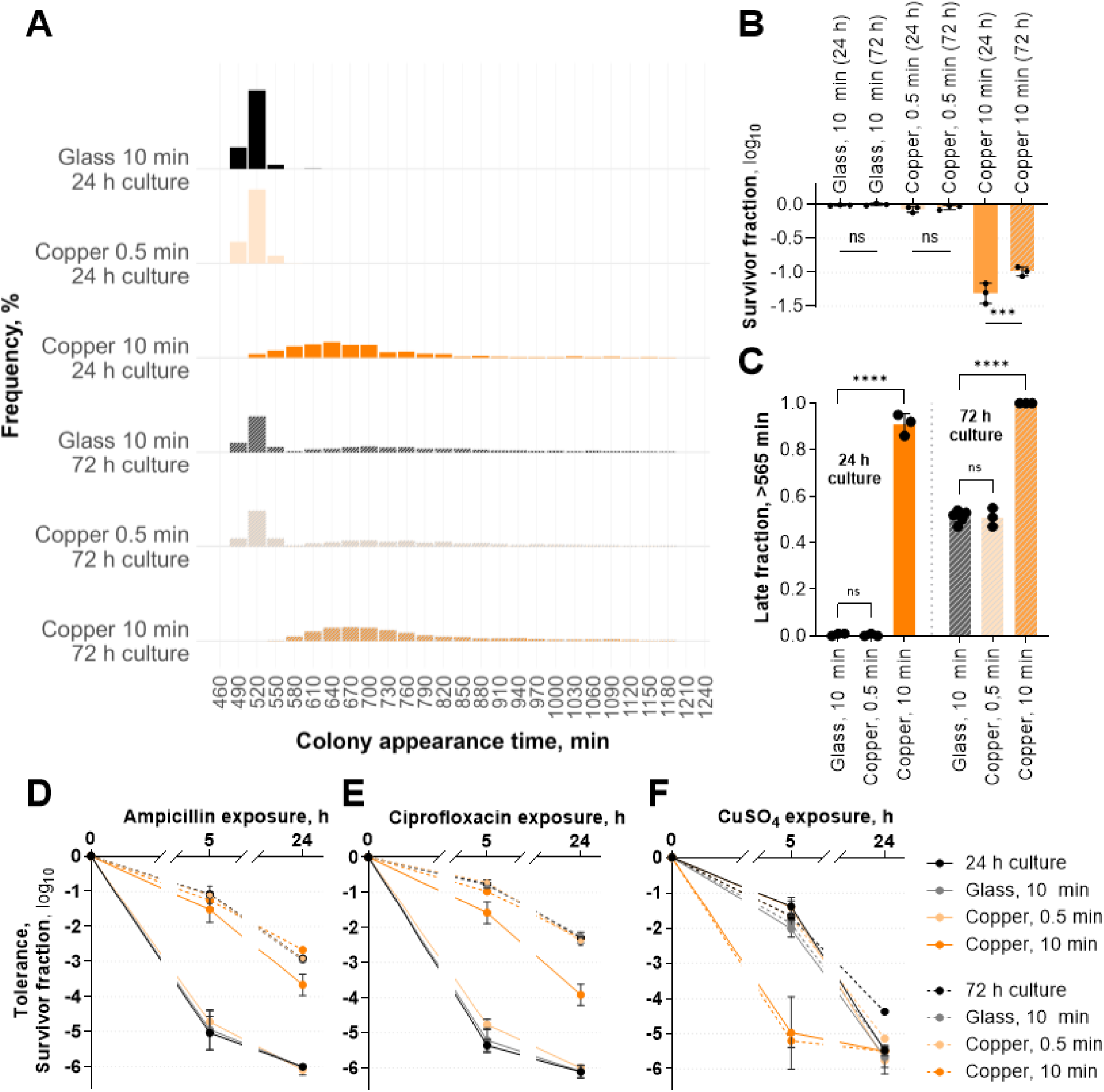
Age of the stationary-phase inoculum affects both copper and antibiotic tolerance. Lag-time distributions (A), post-exposure survival (B), late fraction (C), antibiotic tolerance (D, E), and CuSO₄ tolerance (F) after short biocidal copper surface exposure of inocula prepared from 24 h and 72 h tube cultures. Results from 3 independent cultures and respective exposures ±SD. Statistical comparisons of selected pairs are indicated in panel B with ns P > 0.05, *** P < 0.001, **** P < 0.0001 based on one-way ANOVA with Bonferroni post-hoc testing. Older cultures contained larger pre-existing late fractions and showed higher pre-and post-exposure antibiotic tolerance, consistent with delayed regrowth as a major driver of antibiotic tolerance. In contrast, survivors of copper surface exposure showed reduced tolerance to subsequent CuSO₄ challenge regardless of inoculum lag-tail properties. This divergence indicates that increased antibiotic tolerance after copper exposure is not explained by slower growth alone but is accompanied by copper-specific vulnerability consistent with sublethal injury.

**Supplementary Figure S6.**
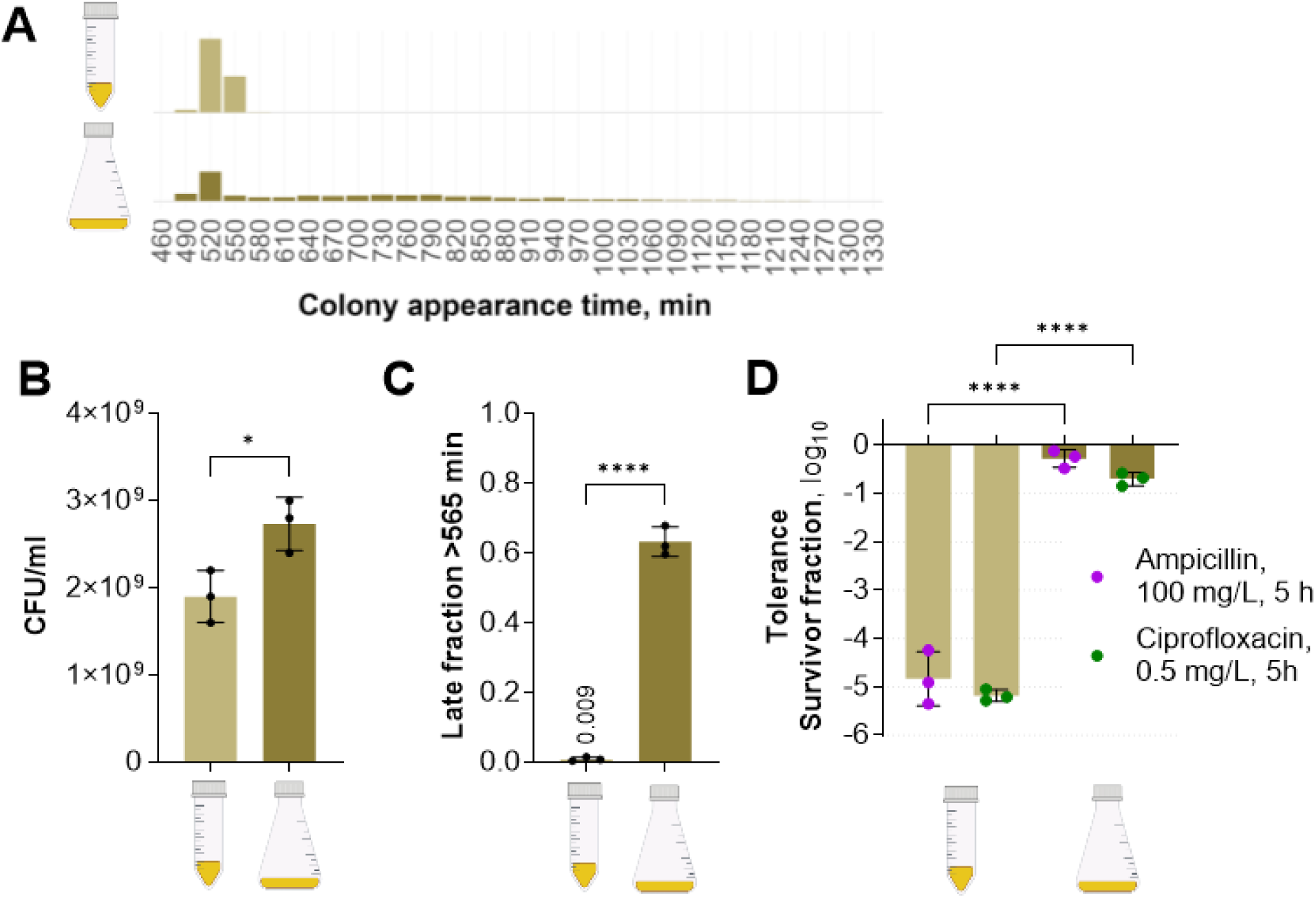
Culture vessel and volume alter baseline late fraction and antibiotic tolerance. Lag-time distributions (A), viable counts (B) and late fraction (C) in cultures and their antibiotic tolerance (D). Cultures were incubated in LB broth at 37°C and 150 rpm for 24 h in tube (10 ml in 50 ml Falcon-type tube at 45° angle, 1/5 volume ratio, not vented) or Erlenmeyer flask (10 ml in non-baffled narrow-neck 100 ml flask, 1/10 volume ratio, not vented). Results from 3 independent cultures ±SD. Asterisks denote statistically significant differences (* P<0.05, **** P<0.0001) based on Students T-test (B, C) or one-way ANOVA with Bonferroni post-hoc testing (C). The tilted tube culture results in 0.86 cm^2^/ml and flask culture in 2.9 cm^2^/ml surface area to volume ratio resulting in ∼3.4x difference that in combination with different vessel geometry possibly results in better aeration of flask cultures and related physiological differences. Culture vessel differences result in higher cell count (B), increased late fraction (C) and antibiotic tolerance (D) of flask cultures compared to same age tube cultures.

**Supplementary Figure S7.**
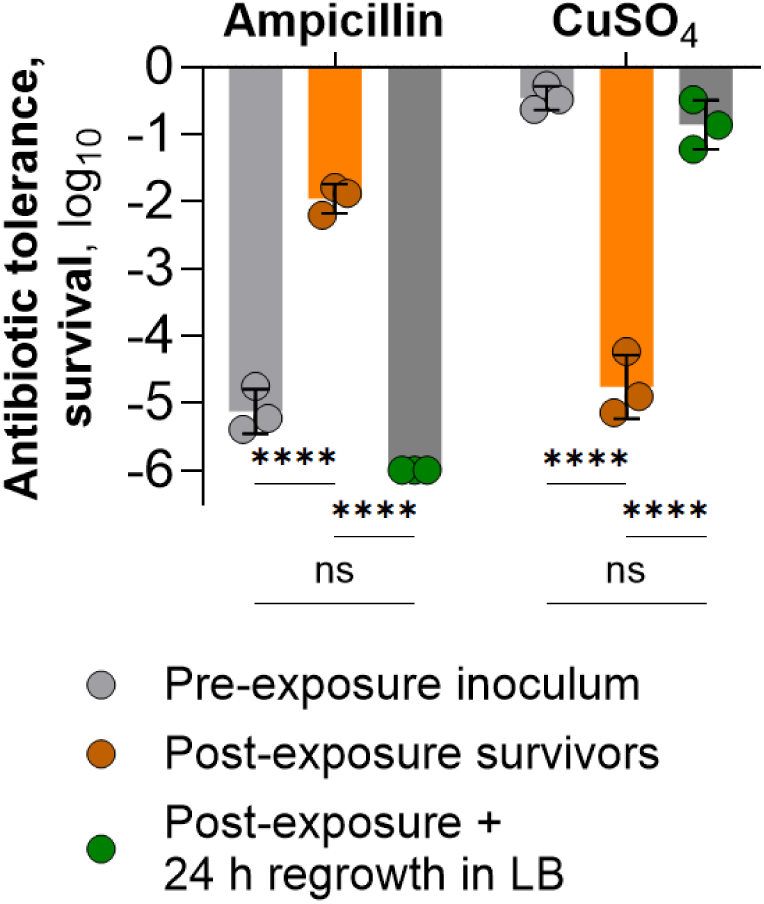
The copper-induced antibiotic-tolerance shift and CuSO_4_ hypersensitivity are transient and reset after regrowth. Ampicillin and CuSO₄ tolerance measured before exposure, immediately after 10 min biocidal copper surface exposure, and after overnight regrowth of the ∼10^6^ CFU/mL survivor population in fresh LB. The copper-exposed population showed increased ampicillin tolerance and decreased CuSO₄ tolerance immediately after exposure, but this phenotype reverted after approximately 10–12 generations of regrowth to stationary phase. Results from 3 independent exposures and respective cultures ±SD. Asterisks indicate significantly different groups by two-way ANOVA with family-wise Bonferroni post-hoc testing (ns denotes P > 0.05 and **** P < 0.0001).

**Supplementary Figure S8.**
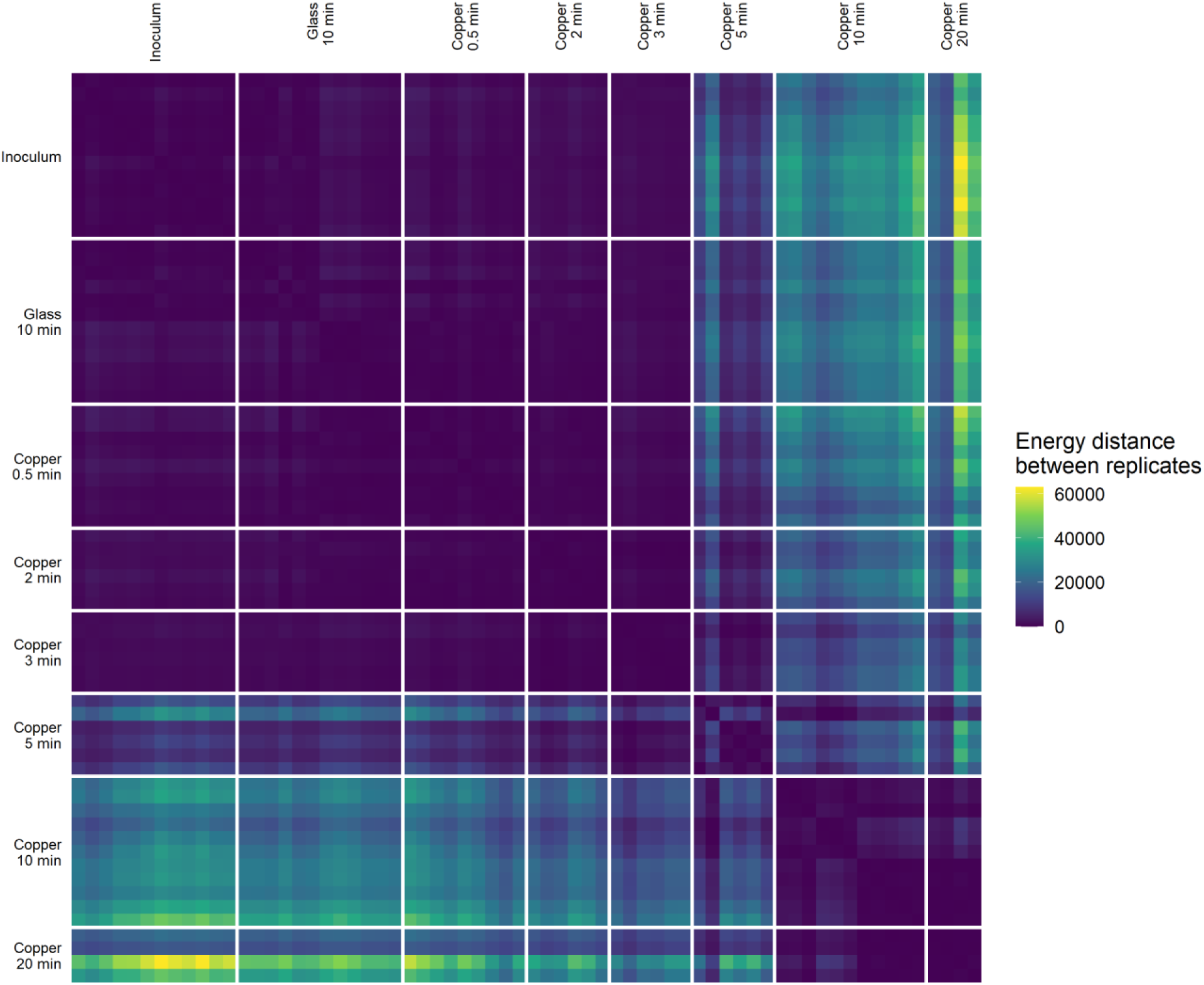
Pairwise energy-distance heatmap of replicate lag-time distributions after incremental semi-dry copper surface exposure. Replicate exposures are ordered by condition, and white separator lines delineate condition blocks. Smaller distances within a block and larger distances between blocks indicate that colony appearance-time distributions are more similar within exposure conditions than between them. Distances were computed at the replicate level from lag-time data used in the main analyses, and the figure complements Fig. 3A by visualizing condition-dependent clustering of post-exposure recovery kinetics across all analyzed conditions.

**Supplementary Figure S9.**
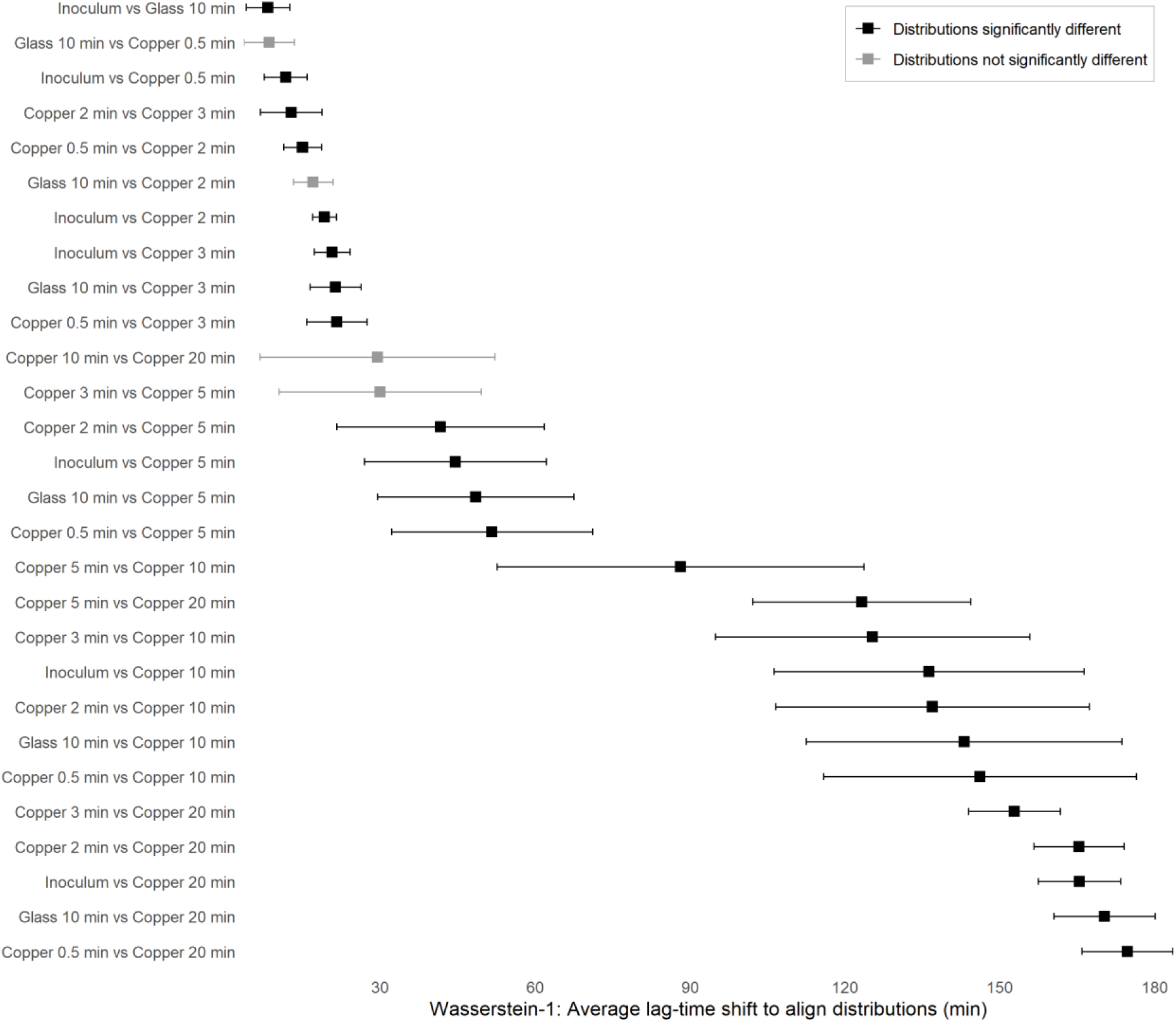
All pairwise Wasserstein-1 (W1) effect sizes for lag-time distributions shifts after incremental semi-dry copper surface exposure. Points show the median W1 across all replicate pairs for each condition pair and horizontal bars denote variability (SD). W1 is expressed in raw minutes and can be interpreted as the average lag-time shift required to align two distributions. Black symbols denote pairs whose lag-time distributions differed significantly by PERMANOVA on energy distances after multiple-testing correction, whereas grey symbols denote non-significant pairs. Larger W1 values therefore indicate progressively greater displacement of recovery-time distributions, especially for comparisons involving longer copper exposures. Figure complements Fig. 3A on which only effect sizes versus 10 min glass reference are presented.

**Supplementary Figure S10.**
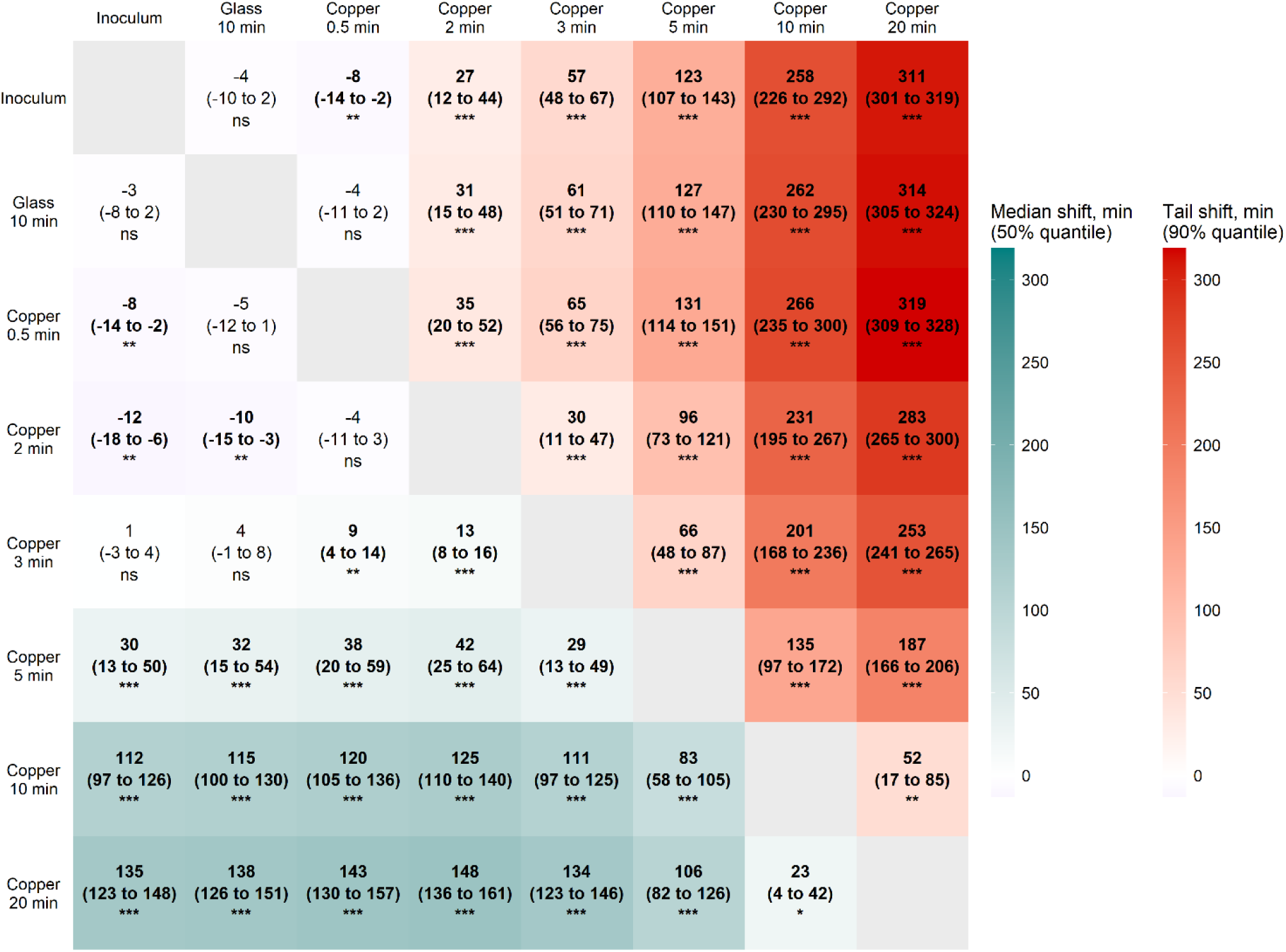
All-to-all lag-time shift matrix after incremental semi-dry copper surface exposure. The lower triangle shows median shift (50% quantile; group2 − group1), and the upper triangle shows tail shift based on the 90% quantile (displayed as group1 − group2 for visual symmetry), both in raw minutes. Cell labels report the estimated shift, 95% bootstrap confidence interval, and adjusted bootstrap significance for the null hypothesis of zero shift; ns denotes P > 0.05, * P < 0.05, ** P < 0.01, *** P < 0.001, and **** P < 0.0001. Positive values indicate later colony appearance in the direction of comparison. **Together, the matrix shows that increasing copper exposure shifts both the center and, more strongly, the right tail of the lag-time distribution to later times.** Figure complements Fig. 3C on which only lag shifts from 10 min glass reference are presented.

